# Maternal characteristics and their relation to early mother-child interaction and cognitive development in toddlers

**DOI:** 10.1101/2024.03.26.586846

**Authors:** Jasmin Preiß, Adelheid Lang, Theresa Hauser, Monika Angerer, Peter Schernhardt, Manuel Schabus

## Abstract

Early mother-infant interaction is believed to have a significant impact on the social, cognitive, and emotional development of children. These interactions are not only influenced by child and contextual factors but also by the mother’s personality traits and strain. In this study, we investigated the relation between maternal factors such as personality, depressive symptoms, or experiencing of emotions, with (i) children’s early cognitive development and (ii) interaction patterns in a sample of 116 mother-child dyads (mean child age = 18.63 months ± 6.42). Maternal factors were assessed using standardized questionnaires, children’s cognitive development was measured using the Bayley Scales of Infant and Toddler Development, and interaction patterns were evaluated using the CARE-Index. The study found that children of mothers who scored higher in agreeableness performed better in cognitive assessments. Additionally, our analysis revealed statistical trends indicating that mothers with higher levels of conscientiousness tended to be less unresponsive in the interaction with their infants, while those with higher levels of neuroticism were more likely to exhibit compulsive behavior in their toddlers. Additionally, there was a trend indicating that maternal depression was associated with increased maternal controlling behavior towards toddlers. Finally, mothers who placed significant importance on their bodily signals to assess their overall well-being had higher scores in the quality of interaction with their child. Overall, these findings show the intricate relation between maternal behavior and state with dyadic interaction quality. This should underline that optimal infant development is only possible if mothers are well supported especially if in need due to various burdens such as depressive symptoms.

## Introduction

### Sensitive parenting behavior

Sensitivity can be defined as the ability to detect and interpret children’s signals accurately and to respond to them appropriately and promptly [1]. Its opposite poles are “unresponsiveness” – where the adult is insensitive to infant stimuli and responds inappropriately – and the dimension “control” where adults are generally aware of infant signals but respond incongruently [2]. Their behavior is typically contingent with the child but can be considered “pseudo-sensitive”, angry or even hostile [2]. Cognitive, social and behavioral delays in development have been associated with emotionally and physically unsupportive family environments in early childhood [1, 3–6], for instance in form of insensitive parenting. In a study on the effect of parental absence, low levels of parent-child interaction have been found to be linked to children’s school and cognitive performance [7]. On the other hand, pleasant emotions – as elicited in an ideal parent-child interaction – are known to have a positive effect on performance in cognitive tasks and creative problem solving [8]. Additionally, academic competence, self-confidence, and positive peer relations have been previously associated with supportive parenting and consistent “discipline practices” [9]. Furthermore, maternal sensitivity and parenting practices have been found to buffer the negative effects of, for example, low socioeconomic status on child outcomes such as cognitive and language performance [10, 11]. These and other findings are consistent with the notion that the quality of early mother-infant interaction is crucial for children’s social, cognitive and emotional development [12, 13]. The far-reaching impact of parents on the development of their children seems to be beyond doubt – but what allows good interaction quality or even drives parental behavior?

### Factors influencing parenting behavior

Amongst various other factors, realistic expectations about the child, parental history of childhood trauma and parental stress are thought to affect parenting [14]. Furthermore, social support has been associated with adaptive parenting, whereas marital problems and divorce, parental illness as well as financial strain have been found to negatively affect parenting capability [15].

A useful framework for understanding the determinants of parenting involves categorizing them into (i) maternal personality, (ii) contextual factors (such as social and emotional support and marital relationship), and (iii) child factors [16, 17]. While maternal reports of child temperament and self-reports of parenting behavior may be inherently prone to biases [18], maternal self-reports remain a suitable method for assessing mothers’ internal constructs such as personality and depression. Consequently, we will focus on maternal (personality) factors in more detail in the following section.

#### Maternal factors

Maternal behavior influences child behavior and vice versa. However, maternal behavior is additionally shaped by prior experiences, expectations and cognitions. The way mothers see their children is thus affected by their own feelings, perceptions and interpretations. Therefore, in order to understand certain parenting behaviors, one has to gain insights into the psychosocial setup of parents. Self-esteem, the subjective evaluation of one’s own worth [19], is discussed to have a significant impact on a mother’s belief to be able to parent her child [20], thus impacting her parenting behaviors. Specifically, low self-esteem, the reflection on negative beliefs about the self, which might be accompanied by dysfunctional assumptions [21], may distort the maternal perception and interpretation of her child’s behavior.

Additionally, prior research has shown that maternal personality traits play a crucial role in shaping parenting behaviors. For instance, Achtergarde et al. [22] emphasized the connection between personality traits and parenting styles in their literature review. Their findings are consistent with a previous meta-analysis conducted by Prinzie and colleagues [23], finding favorable parenting practices to correlate with high agreeableness, openness to experience, and conscientiousness, and lower levels of neuroticism. Conversely, higher levels of maternal neuroticism were associated with tendencies toward power-based parenting, reduced support for their children, and an increase likelihood of reporting behavioral problems [22]. Furthermore, Prinzie et al. [23] found lower levels of neuroticism to be linked with “warm” parenting. In a sample of Japanese mothers, Jing and Michiyo [24] identified an association between higher neuroticism and dismissing and dysfunctional parenting responses to child anger, as well as a link between higher maternal conscientiousness and reduced noninvolvement. Besides, a study by Bailes and Leerkes [25] revealed that maternal agreeableness predicted a decrease in negative maternal behaviors.

As one of the key tasks of the mother is the regulation of an infant’s affect it seems reasonable that maternal experience of her own emotions, meaning the internal process of perceiving, understanding, and managing emotions, as well as her strategies to deal with her feelings have an impact on the interaction with the child. Meyer and colleagues [26] for example found that 4–5-year-old children of parents with high emotion suppression are less likely encouraged to express their emotions. Furthermore, preoccupation with negative emotions or depression is thought to impede the mother to pay attention to her child’s signals or to be emotionally available. Regarding postnatal depressive symptoms many studies indeed find lower maternal sensitivity [27, 28] associated with depressive symptoms. A systematic review by Deans [29] however suggests the relationship between maternal depression and sensitivity to be more complex. While depression can pose a risk factor for reduced sensitivity, its impact is not universally disruptive. As cited in Deans, multiple studies have showcased varying outcomes, highlighting instances where depressed mothers displayed similar sensitivity levels to non-depressed counterparts [30–33]. However, other studies did find some impacts of depression. For example, maternal depression together with poverty can notably impact maternal sensitivity [34, 35]. Furthermore, in their study, Hummel and colleagues [36] discovered that mothers, regardless of depressive symptoms, exhibited comparable positive affect when interacting with their toddlers. However, mothers experiencing moderate to high levels of depressive symptoms exhibited a lack of reciprocal positive engagements with their two-year-olds, a contrast to the interactions observed among other mothers.

### Study Aim

Our study aims to further explore how maternal factors influence interactions with their children, focusing on infants (birth to 15 months) or toddlers (16 to 35 months). We seek to analyze the complex associations between maternal factors such as personality, depressive symptoms, self-esteem, and emotional experiences, and interaction patterns between mothers and their children. Utilizing direct observation, we employed Crittenden’s Child-Adult Relationship Experimental Index (CARE-Index; [2]), a well-established tool for comprehensive assessment of dyadic interaction quality and behavioral patterns. This method allows for objective evaluation of external behaviors and interactions. Additionally, we aim to explore the predictive value of these maternal factors on the cognitive development of the child.

## Materials and methods

### Participants

Overall, a total of 110 German-speaking mother-child dyads were assessed and recorded using videography. Based on the applied tool to assess interaction (i.e., the CARE-Index; cf. chapter 2.3.1), the sample was split into two groups: Mother-infant dyads *(n* = 38; assessed with the Infant CARE-Index) and mother-toddler dyads (*n* = 72; assessed with the Toddler CARE-Index).

In the 38 Infant CARE-Index interactions, the mothers were on average 31.34 years old (*SD* = 4.98; range = 17 - 45) and the infants (19 being female) were on average 11.55 months old (*SD* = 2.34; range = 6 - 16). In the 72 Toddler CARE-Index interactions, the mothers were on average 31.63 years old (*SD* = 5.13, range = 21 - 44) and the toddlers (39 being females) were on average 21.85 months old (*SD* = 4.46; range = 15-35). At the time when cognitive development was assessed with the Bayley Scales of Infant and Toddler Development, toddlers (*n* = 41) were on average 25.49 months old (*SD* = 2.30; range = 22-32). The mean time span between cognitive assessment and CARE-Index assessment was 2.54 months (*SD* = 3.13; range = 0 - 10.87).

In our sample, 46% of the mothers had a university degree, indicating a relatively high level of educational attainment among the participants, and 96 % were in a partnership or relationship. The majority of mothers, comprising 93% of the total, held citizenship from either Austria (77%) or Germany (16%). The mothers gave written informed consent and participated on a voluntary basis. For the minor mother involved in our study, her legal guardians provided supplementary written consent.

The presented study was approved by the ethics committee of the University of Salzburg (EK-GZ 12/2013). Recruitment occurred between April 10, 2017 and February 21, 2020.

### Procedure

The cross-sectional experimental design of the study is illustrated in *Fig 1*. Participants were asked to come for a recording session to the “Laboratory for Sleep, Cognition and Consciousness Research” in Salzburg, Austria. There, the assessment of the mother-child interaction was conducted. Depending on the children’s age and emotional status, their cognitive development was being tested subsequently or at a separate appointment. Thereafter, the mothers received an online link for self-report questionnaires including the Big-Five Inventory-10 (BFI-10 [37]), Beck Depression Inventory-II (BDI-II [38]), Multidimensional Self-Esteem Scale (MSWS [39]), and a questionnaire on the experience of emotions (Skalen zum Erleben von Emotionen; SEE [40]).

**Fig 1.**
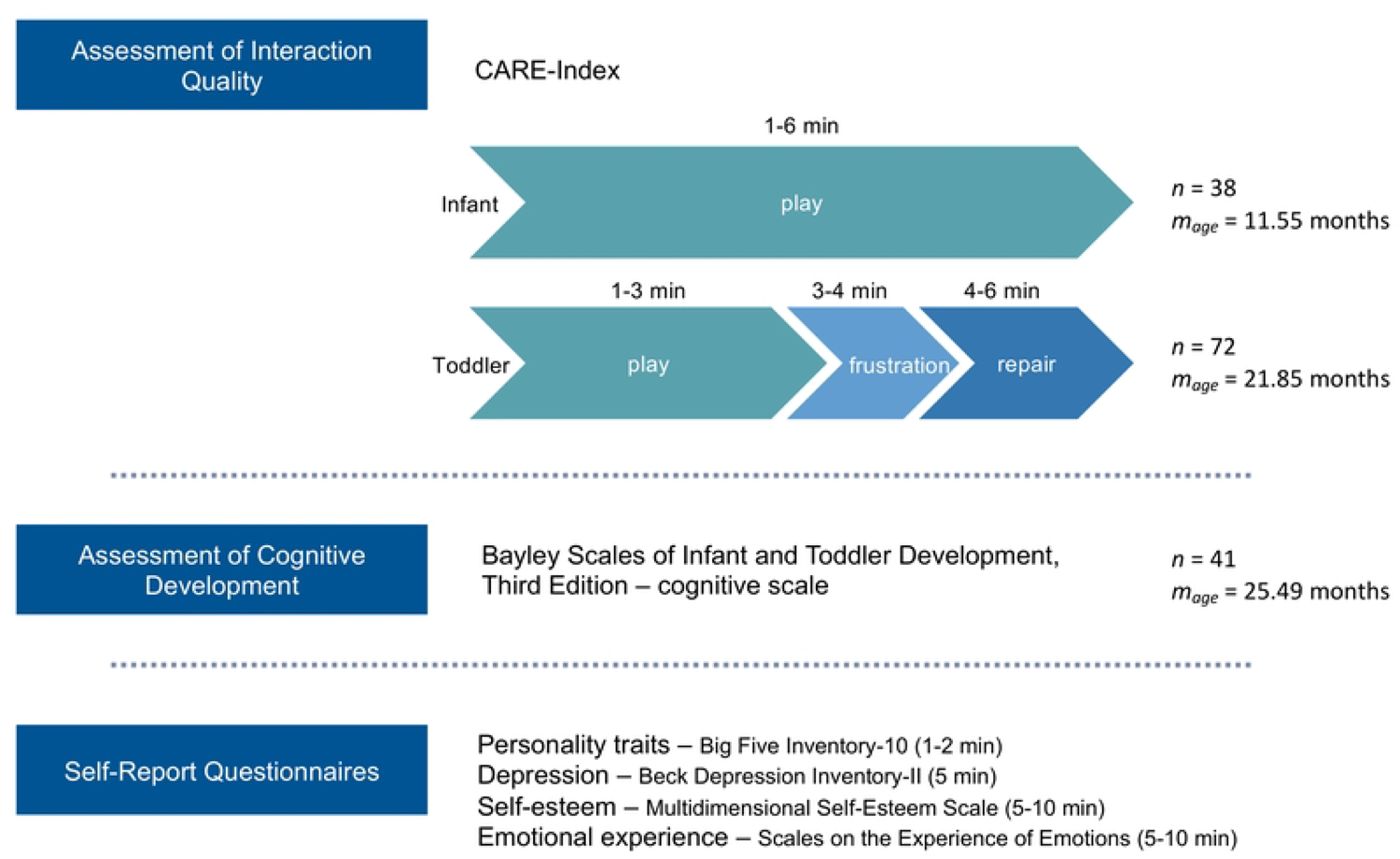
Experimental Design. Mother-child interaction was assessed with the Child-Adult-Relationship Experimental Index (CARE-Index, Crittenden, 2010). The setup of the CARE-Index is displayed separately for infants (up to 15 months) and toddlers (15 months and older). The numbers specify the duration of the different parts (play, frustration, and repair) of the CARE-Index procedure. Toddler’s cognitive development was assessed with the Bayley Scales of Infant and Toddler Development. Self-report questionnaires were used to assess maternal personality, depressive symptoms, self-esteem, and emotional experience.

### Measures

#### Mother-child interaction

The quality of the mother-child interaction was measured using the CARE-Index (CI [2]). It is based on a videotaped adult-child playful interaction for which a variety of age-appropriate and age-inappropriate toys is provided. Parents are allowed to choose which toys to use and may also not use toys at all. The Infant CARE-Index (ICI) functions on the basis of the instruction: *“Play with your child as you usually would.”* And is applied in the context of infants (15 months of age or less). The Toddler CARE-Index (TCI) is used for toddlers (15 months or older), and includes a “frustration-task”, where, after three minutes of play-time, the caregiver is signaled to frustrate the child, for instance by breaking the rules of a game. After one minute, a repair-phase, during which one should return to a play in a way that it makes the child comfortable again, follows. The caregiver receives the additional instructions regarding the phases prior to the assessment. In this study, a fixed six-minute time window for the interaction has been determined to ensure standardization and emphasize the repair phase, which is considered to be of great informational value. The timings are based on recommendations of CARE-Index experts.

Seven aspects of interactional behavior (facial expression; vocal expression; position and body contact; expression of affection; turn-taking; control of the activity; developmental appropriateness of the activity) are rated separately for adults and children. The CARE-Index differs from several observational methods tallying the frequency of specific behaviors. Instead, it prioritizes observing interaction patterns and behavior is scored by its function and based on the perspective of the interaction partner (for more detail, please see [41]). Adult scales are “sensitive”, “unresponsive” and “controlling”. Infant (birth – 15 months) scales are “cooperative”, “compulsive”, “difficult” and “passive”, while toddlers (15 – 36 months) are rated “cooperative”, “compulsive”, “threateningly coercive” and “disarmingly coercive”. Dyadic synchrony (DS) combines caregiver sensitivity with child cooperation and is defined as the fit between adult and child. Scores on all scales range from 0 to 14, with higher scores indicating more observations of the construct. In terms of the DS, a score of 11-14 represents sensitive, 7-10 adequate, 5-6 inept interaction, while 0-4 indicates high risk child development.

Each video recording was rated by at least one reliable rater, with 30% of the videos being additionally coded by another independent rater. The interrater intraclass correlation coefficient (ICC) for the dyadic synchrony of eleven mother-infant interactions was 0.87, indicating good reliability. For the 23 randomly selected mother-toddler interactions, a comparably good reliability with an ICC of 0.79 was achieved. For more details on each of the subscales please see *S1 Supporting information, Suppl. Table 1*.

#### Maternal factors

The 10-item Big Five Inventory (BFI-10 [37]) was used to measure the mothers’ *personality features*, based on the Big Five dimensions: openness to experience, agreeableness, extraversion, neuroticism, conscientiousness. Each dimension is rated on a 5-point Likert scale ranging from 1 (totally disagree) to 5 (totally agree). The BFI-10 scales, on average, accounted for 70% of the total variance observed in the full BFI-44 [42]. Additionally, they maintained an 85% retention of retest reliability.

Maternal *depressive symptoms* were measured with the German version of the Beck Depression Inventory-II (BDI-II [38]), a 21-item self-administered questionnaire with a high internal consistency (.84 ≤ a ≥ .91 in non-clinical samples [38]).

A questionnaire on the experience of emotions (Skalen zum Erleben von Emotionen; SEE [40]), which is based on Carl Roger’s client-centered personality theory, was utilized to assess how the mothers *experience, appraise and deal with their emotions*. The 42 items are subdivided into seven independent scales: *acceptance* of own emotions, experienced emotional *overflow*, experienced *lack* of emotions, body-related symbolization of emotions (*somatization*), imaginative symbolization of emotions (*imagination*), experience of emotion *regulation*, experience of *self-control.* All scales demonstrate internal consistency within the range of alphas from .70 to .86 [40].

Maternal *self-esteem* was assessed with the Multidimensional Self-Esteem Scale (“Multidimensionale Selbstwertskala”; MSWS [39]), an adaptation of the Multidimensional Self-Concept Scale by Fleming and Courtney [43], which comprises many facets of the concept self-esteem. For this study, the focus was placed on global self-esteem (“MSWS global”), which is based on all six subscales (emotional, social [social contact, handling criticism], competence, physical attractiveness, sportiness). The higher-order scales exhibit strong internal consistency, with alpha values ranging between .84 and .90.

#### Child cognitive development

To assess the cognitive development, the cognitive scale of the Bayley Scales of Infant and Toddler Development, Third Edition – German Version (Bayley-III [44]) was administered by a trained female examiner in the presence of the mother. Children either sat on their mother’s lap or on their own chair at a table. The cognitive scale includes items to examine e.g. the sensorimotor development, exploratory and manipulative behavior, object-centeredness and memory [45]. The 91 items are organized in ascending order of difficulty and the item raw scores (total number of correctly solved items) are being transformed into age-normed sub-scale scores (ranging from 1-19, with a mean score of 10 and a standard deviation of 3). To enable comparison with the motor and language scales, scale-value-equivalents (ranging from 40 to 160 with a mean of 100 and a standard deviation of 15) can be calculated.

#### Data analyses

Means and standard deviations are reported for the description of the samples. Non-parametric Spearman’s correlations were used for the bivariate analyses of the ordinal variables. The significance level was set to α = 0.05 (two-sided) for all analyses. To adjust for multiple comparisons, FDR (False Discovery Rate) was applied and additionally reported. As suggested by Wasserstein and colleagues [46], we interpreted the overall pattern rather than focusing on individual *p*-values. Therefore, we also interpreted *p*-values 0.05 < *p* ≤ 0.10 if they were in line with the overall pattern of results and if the *p*-values remained within this range after correction. Statistical analyses were conducted using IMB SPSS Statistics, Version 27.0 and RStudio 2023.12.0 were used for statistical analyses.

## Results

### Mother-child interaction

In the infant sample, a positive correlation between the CARE-Index subscales maternal control and children’s compulsiveness was found (*r_s_* = .50, *p* = .001, *p_corrected_* = .004) indicating that more controlling mothers also tend to have more compulsive infants. This association was however not significant in the toddler sample (*r_s_* = .19, *p* = .110, *p_corrected_* = .147), indicating that the coherence between increased maternal control and elevated compulsive behavior is in our sample limited to infants. For a detailed overview on the CARE-Index scores, as well as the correlations between the scales see *S1 Supporting information, Suppl. Table 2*.

### Maternal Characteristics and Interaction Quality

#### Maternal Personality Traits

Spearman rank correlations were performed in order to test for associations between maternal personality traits (BFI) and mother-child interaction patterns.

As illustrated in *Fig 2 A*, maternal conscientiousness was found to be negatively correlated with unresponsive maternal behavior (*r_s_* = -.41, *p* = .016, *p_corrected_* = .062), indicating a trend toward statistical significance. This suggests that mothers who are more conscientious tended to be more responsive towards their infants (*n* = 34). In the older toddler sample (*n* = 61), higher scores of maternal neuroticism (BFI-10) were found to correlate with children’s display of more compulsive behavior (*r_s_* = .31, *p* = .017, *p_corrected_* = .067, cf. *Fig 2 B*), also indicating a trend towards significance. For more details, please see *S1 Supporting information, Suppl. Table 3*.

**Fig 2.**
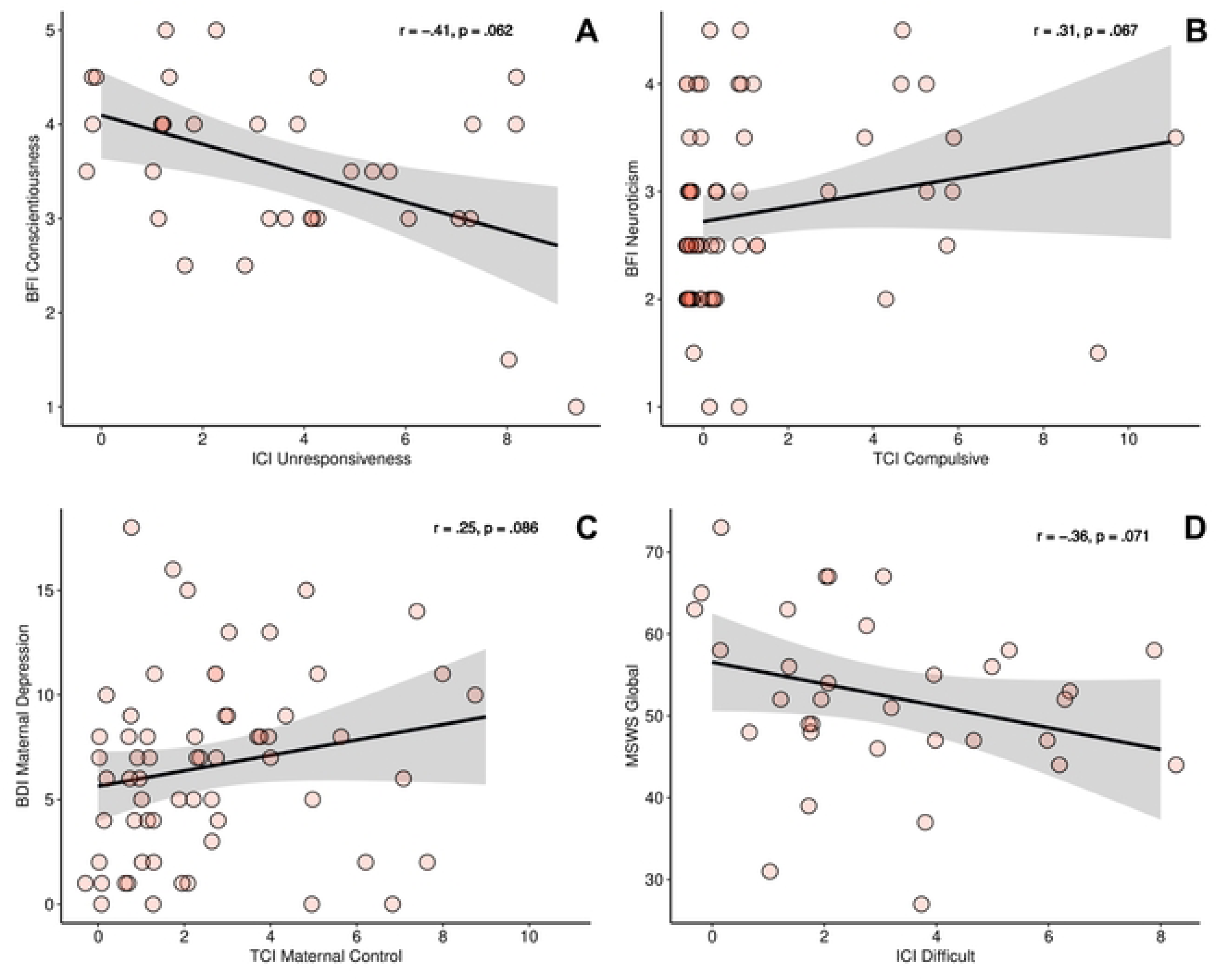
Maternal characteristics and mother-child interaction. **(A)** In mothers of infants (*n* = 34), there is a trend suggesting that higher maternal conscientiousness scores are associated with lower unresponsive maternal behavior. **(B)** A trend indicates higher maternal neuroticism scores to be linked with more display of compulsive behavior in toddlers (*n* = 61). **(C)** Among mothers of toddlers (*n* = 62), there was a trend indicating that higher maternal depression scores were correlated with more maternal controlling behavior. **(D)** Additionally, a trend was observed suggesting that lower overall self-esteem (MSWS Global) was associated with more difficult behavior in infants (*n* = 35). Abbreviations: BFI = Big Five Inventory; ICI = Infant CARE-Index; TCI = Toddler CARE-Index. BDI = Beck Depression Inventory; MSWS = Multidimensional Self-Esteem Scale. The shaded area in grey represents the standard deviation.

#### Maternal depression and self-esteem

Although maternal depression, as indicated by a mean Beck Depression Inventory-II (BDI-II) score of 6.53 (*SD* = 4.37), was subclinical on average among mothers of toddlers (*n* = 62), our analysis revealed a trend suggesting that higher maternal depression scores were associated with increased maternal controlling behavior in this sample (*r_s_* = .25, *p* = .048, *p_corrected_*= .086, see *Fig 2 C*).

Similarly, infants (*n* = 34) of mothers with an overall lower self-esteem (MSWS global) displayed more difficult behavior during the CARE-Index interaction: *r_s_* = -.36, *p* = .040, *p_corrected_* = .071, indicating a trend (cf. *Fig 2 D*). For more details, please see *S1 Supporting information, Suppl. Table 4*.

#### Maternal experience of emotions

In mothers of infants (*n* = 25) statistical trends suggested a positive correlation between higher scores in the SEE subscale “somatization” and higher dyadic synchrony (*r_s_* = .46, *p* = .020, *p_corrected_* = .097, see *Fig 3 A*). Additionally, there was a trend indicating that higher sub-scores for infant cooperation were associated with higher somatization scores (*r_s_*= .47, *p* = .019, *p_corrected_* = .095) suggesting that mothers who apprehend their bodily sensations to mirror their emotional state may have more cooperative infants, potentially contributing to higher quality interactions. For more details, please see *S1 Supporting information, Suppl. Table 5*.

**Fig 3.**
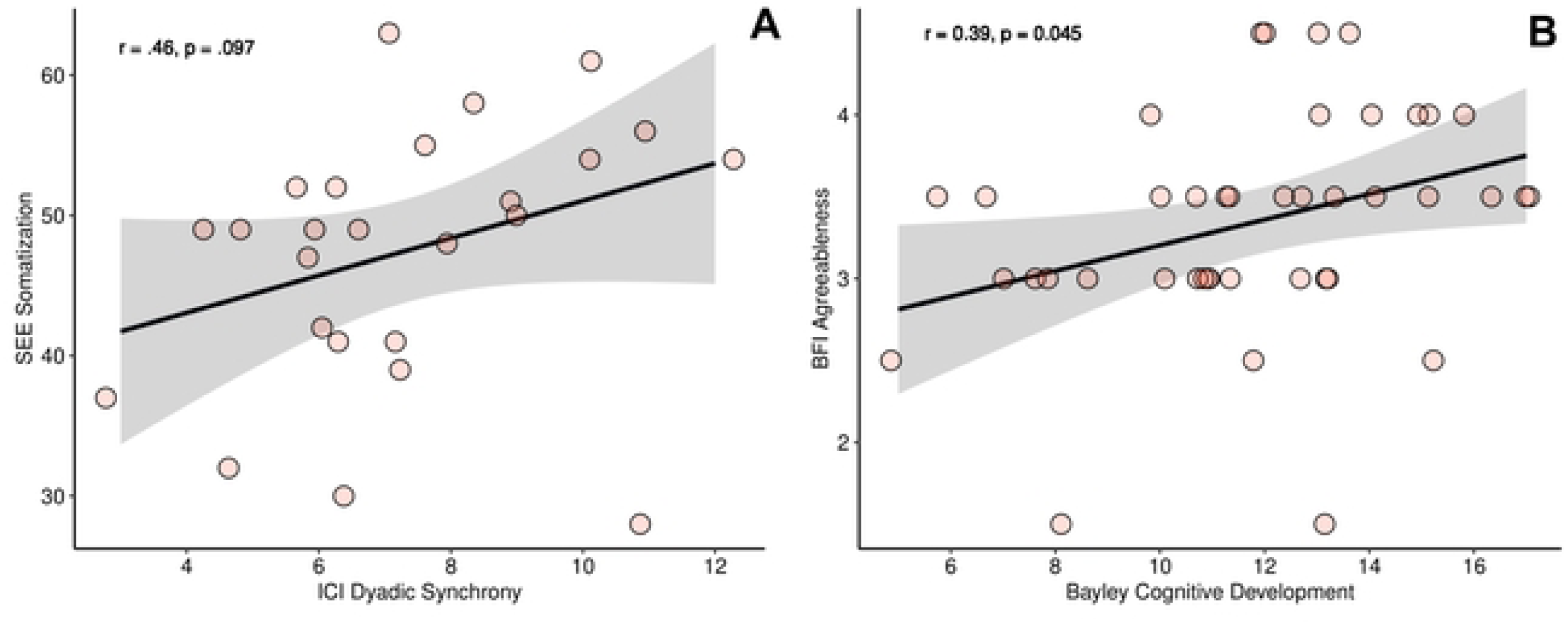
Maternal characteristics, mother-child interaction and toddler cognitive development. **(A)** A trend is suggesting that mothers who are more attuned to their bodily signals (higher “somatization” in SEE) exhibit higher dyadic interaction quality with their child (*n* = 25). **(B)** Higher maternal agreeableness is associated with superior cognitive development in toddlers (*n* = 41) with cognitive development being around the norm (range 8-12) at that age. Abbreviations: SEE = questionnaire on the experience of emotions; ICI = Infant CARE-Index. The shaded area in grey represents the standard deviation.

#### Maternal factors and child cognitive development

The cognitive development of 41 toddlers was assessed with the Bayley scales. Children’s cognitive development scores ranged from 5 to 17 with a mean score of 11.83 (*SD* = 2.95) that lies within the norm range (8-12).

We found that mothers with higher agreeableness scores have children with superior performance in the Bayley scales of cognitive development (*r_s_* = .39, *p* = .013, *p_corrected_* = .045, see *Fig 3 B*). For more details, please see *S1 Supporting information, Suppl. Table 6*.

## Discussion

In the present study, we focused on maternal characteristics including e.g., personality and self-esteem, but also maternal strain such as depression as well as general emotional experiencing and their effect on mother-child interactions (using the CARE-Index), and on infant’s cognitive development.

One of the main findings was that maternal agreeableness was positively associated with cognitive development in the toddler suggesting this maternal quality to support early child development. Mothers who use their body to interpret the meaning of their emotions interestingly tended to show interaction patterns with higher dyadic synchrony, indicating maternal self-awareness to be beneficial for the interaction with the child. Additionally, we found some age-specific (i.e., infants, toddlers) relations between child behavioral patterns and maternal personality traits.

### Maternal personality

Looking at maternal personality, we found a potential link between unresponsive maternal behavior and personality traits: In mothers of infants, higher levels of unresponsiveness in the play interaction were correlated with less conscientiousness. Unresponsive maternal behavior can be generally described as being more withdrawn or less attentive to children’s signals and needs. According to [47] the primary function of such maternal behavior is the reduction of contact or engagement, and these interactions lack structure and goal-orientation in the play with the child. Conversely, conscientiousness is associated with being organized, responsible, reliable, goal-directed, hardworking, self-disciplined and thoughtful [48–50]. While conscientiousness has been previously linked with responsive maternal behavior – a concept that is comparable to the DMM’s term “sensitivity” [48] – we found that more conscientious mothers were less unresponsive but not necessarily more sensitive. Instead, our results indicate that more conscientious mothers tend to be more controlling or sensitive in the interaction with their child. It is possible that more conscientious people feel obliged to interact more actively, potentially also taking control or imposing their will in play situations which may sometimes be a bit too much for the child. One possible explanation for these similar but somewhat different findings could be the role of structure: both, responsiveness and control are associated with a certain amount of structure and predictability. In accordance with this, Prinzie and colleagues [23] suggested conscientious parents to establish their “high standards” in child rearing and thereby offering a more ordered setting. However, too much conscientiousness might even lead to a structure that is too rigid and not adaptive to the situation and therefore would come across as too “controlling”. This could be in line with the findings of Karreman et al. [51]. In their study they differentiated between positive control, which offers structure, limits and clarifications, and negative control, functioning on the exertion of control through punitive and harsh parenting methods. Both forms of control had different moderating effects between parenting and problem behavior in 36-month–old children [51].

A statistical trend suggested a potential link between greater maternal neuroticism and with toddlers’ more pronounced display of compulsive behaviors. In other words, mothers with a predisposition to experience intense negative emotions or a lack of emotional stability [52] were associated with children who had learned to inhibit their negative affect and to substitute their actual affect with a false positive one. Foundation of the co-occurrence of this maternal propensity and their toddler’s behavior might be the mother’s prevalent impression of an unpredictable and uncontrollable environment [53] that feels unsafe for her and her offspring causing her to be overprotective. In accordance with this, Clark and colleagues [48] found higher levels of neuroticism to be linked with intrusive parenting behaviors. Intrusiveness is commonly associated with parenting behavior in the context of children’s compulsiveness and has even been suggested as a possible predictor of child maltreatment [54]. However, it is important to note that the inter-rater reliability, as measured by the intraclass correlation (ICC), was found to be low for toddlers’ compulsive behavior. This may be due to a less clear pattern of compulsive behaviors in toddlers, or it may indicate that the scale is not well-suited for this age group, suggesting a potential need for modification. Therefore, interpreting this result requires caution, highlighting the need for additional research to gain a more comprehensive understanding of toddlers’ compulsive behavior and thoroughly investigate the relationship between maternal neuroticism and toddlers’ compulsiveness.

### Infants vs. toddlers: The role of development

As outlined above, some of our findings were age specific (i.e., only visible in infant or toddler interactions). It is plausible that different developmental tasks need to be accomplished depending on children’s maturational levels, prompting dynamic and versatile caregiving behaviors in order to maintain sensitivity and adaptability. Besides that, information processing becomes increasingly complex through maturation and the development of a wider range of more complex strategies of the child becomes possible [47]. Furthermore, young infants’ patterns might be less pronounced, which would also emphasize the importance of early interventions or prevention, whereas in toddlers the dyadic dysfunction may already have aggravated over time and is thus more stable.

### Strain and Maternal Emotions

Focusing on further variables on the side of the mother, one might assume a connection between depressive maternal symptoms and unresponsive dyadic behavior with the child, reflecting their disengagement due to lower levels of positive mood. However surprisingly, we did not find higher maternal depression scores to be related to unresponsiveness. Rather a trend suggested maternal depression to be associated with more controlling behavior during the dyadic interaction with their toddlers.

This seems in line with a meta-analysis by Lovejoy et al. [55] on maternal depression and parenting behavior. Lovejoy and colleagues found a strong relation between depression and hostile maternal behavior, as well as a “less powerful relationship with disengagement”. Humphreys and colleagues [56] analyzed speech samples of mothers of six-month-old children based on narratives. They found that the use of language of mothers with higher depressive symptoms often comes with greater psychological distance and higher levels of self-focus, which in turn can be associated with less caregiver warmth [56]. Overall, this could be seen as in accordance with controlling behavior, that is more “harsh” parenting and the adult’s perception of the children’s signals without responding adequately and rather following their own plan. Especially for mothers with depressive symptoms that may be the only way to engage in the first place as they have less capability to identify the needs of the child adequately and sensitively as they are occupied with their own negative feelings.

Further, we found that mothers who have a more pronounced belief that their bodily perceptions are meaningful and are associated to their mental state tend to have a higher interaction quality with their (more cooperative) infants. This supports the notion that interoception, the competence to be aware of internal bodily changes [57, 58], is of great importance for caregiving [59]. One of the core responsibilities of new parents is co-regulation. Through co-regulating their infant, caregivers give them an idea of what they physically and emotionally feel by responding to their signals timely and appropriately, which in turn provides the base for development of a child’s sense of self and self-regulation. Thus, if a caregiver is well aware of its own internal bodily processes, one can assume that it is easier to identify the child’s needs and to engage in interactions that are well attuned. Important however is the balance: A mother who is fully aware of and focused on her infant’s (physiological) needs but does not acknowledge her own will not be able to support her child properly either in the long run. Hence, it seems like there is a need of a balance between the perception of and response to the children’s and the own needs that may change over time from a stronger focus on children’s needs that is more equally distributed as the child matures.

### Maternal self-esteem

An additional statistical trend suggested lower maternal self-esteem to be related to more display of infant difficult behaviors such as grimacing, vocal protests or refusal to engage in playful dyadic interaction. Related to this, Leerkes and Crockenberg [20] found maternal self-efficacy, which is thought to predict self-esteem, to be linked to the ability of infants to calm down. It is speculated that, also in our study, mothers with higher self-esteem are believing more in their skills to handle or sooth their child, which provides a safer environment for the child and therefore less prevalent difficult child behaviors in interaction situations.

### Maternal factors and child cognitive development

Finally, we did find maternal personality to be linked with toddlers’ cognitive development. Superior performance in the assessment of toddlers’ cognitive development was positively correlated with higher maternal agreeableness, a personality trait, which is not only linked to kindness and warmth, but also to enhanced self-regulation capabilities [60] and (perceived) emotional support of others [61]. Together with our finding of a trend towards a correlation between higher agreeableness and a better interaction quality, one can assume that these mothers might better succeed in sensing and supporting their children’s emotion (regulation) and maintaining a regulated state, enabling the children to explore and interact in the new assessment environment. Yet, further research is needed to elucidate the directionality of this relationship.

The implementation of the CARE-Index, which is an objective measurement of interaction patterns and interaction quality, is what we consider a great strength of our study design. In our view it gives considerably more insight into dyadic patterns than simple self-report questionnaire data.

### Limitations

The study has several limitations that require acknowledgment. Firstly, the sample size was relatively small, and the inclusion of broad age ranges in both groups may have introduced variability.

It is important to note that the identified effects are correlational in nature. The interpretations provided are based on plausible explanations derived from existing literature and our understanding of attachment theory. Unfortunately, the study lacks longitudinal data, which would have allowed us to track the developmental trajectories of these infants and toddlers over subsequent years. We anticipate that future research endeavors will shed more light on these trajectories.

Another potential limitation may arise from the mean time of 2.5 months between the evaluation of dyadic interaction and cognitive development assessment. While maternal sensitivity has demonstrated relative stability across early childhood stages [62, 63], cognitive development tends to adapt swiftly. However, even after controlling for the duration between assessments, the results remained unchanged.

Finally, this study did not include information on paternal characteristics or father-child interactions, nor did we measure the amount of time parents spent with their children. In our upcoming longitudinal study, inspired by the insights gained here, we aim to address these methodological gaps by conducting a comprehensive perinatal study that includes fathers and father figures.

## Conclusions

To summarize, we found several maternal trait factors such as neuroticism, agreeableness, or how a mother is dealing with her own personal feelings that were associated with mother-child interaction patterns. Particularly, our findings highlight the importance of professional support for mothers who experience psychological stress or have unfavorable predispositions such as low self-esteem, depression, or high levels of neuroticism due to their own attachment and parenting history. Providing more accessible parental assistance and support early in development could enhance the quality of interaction and attachment, leading to a healthier and more self-sufficient future generation.

## Acknowledgments

The authors wish to thank the participating mothers and children for their willingness to take part in this project as well as the students who helped with data collection.

## S1 Supporting information

**Suppl. Table 1.** Intraclass correlation coefficients (ICC) for the CARE-Index subscales for infants (n = 11) and toddlers (n = 23)

**Suppl. Table 2.** CARE-Index interaction quality: Means, standard deviations, and spearman’s correlations for infants (n = 38) and toddlers (n = 72)

**Suppl. Table 3**. Maternal personality traits (BFI-10) and mother-child interaction: Means, standard deviations, and spearman’s correlations split for infants (n = 34) and toddlers (n = 61)

**Suppl. Table 4.** (A) Maternal depression (BDI-II) and mother-child interaction: Means, standard deviations, and spearman’s correlations in toddlers (n = 62). (B) Maternal experience of emotions (SEE) and mother-child interaction: Means, standard deviations, and spearman’s correlations for infants (n = 25)

**Suppl. Table 5.** Maternal self-concept (MSWS) and mother-child interaction: Means, standard deviations, and spearman’s correlations for infants (n = 34) and toddlers (n = 61)

**Suppl. Table 6.** Child cognitive development (Bayley) and maternal factors: Means, standard deviations, and correlations with confidence intervals in the toddler sample (n = 41)

